# Negative resilience trends to more frequent extreme droughts in Mediterranean oak species

**DOI:** 10.1101/2024.11.19.624310

**Authors:** I. J. Borreguero, Á. Sánchez-Miranda, E. Martínez-Sancho, A. E. Rubio-Casal, R. Sánchez-Salguero, L. Matías

## Abstract

Recent changes in climate have triggered widespread mortality events in oak forests worldwide. Despite the ecological importance of these ecosystems, limited information exists on the long-term resilience of oak forests in response to extreme climate events. *Quercus canariensis*, a semi-deciduous oak sensitive to summer drought characteristic of the Mediterranean climate, is currently showing defoliation and mortality episodes in the drier areas of its natural range. Here, we investigated the long-term impacts of climate on the radial growth of *Q. canariensis* and assessed changes in resilience to extreme droughts across eight populations in S Spain following an environmental gradient. We observed a clear latitudinal gradient in growth performance, as reflected by resilience indices across populations. Since 1970, *Q. canariensis* has shown a marked decline in resilience to extreme drought events at the wettest region, resulting in a diminished capacity to recover pre-drought growth levels. This trend has preceded growth declines in certain populations, suggesting increased vulnerability to the increasing frequency of extreme climatic events. Through growth analyses, we identified early-warning signals of forest decline in this Mediterranean oak species, highlighting the most sensitive populations and its high susceptibility to extreme droughts impacts.

## Introduction

The observed changes in climate over the last decades have modified the phenology, growth, and biotic interactions of plant species across the globe (Menzel et al., 2020; Parmesan, 2006). The Mediterranean region has been identified as a climate-change hotspot, due to its sensitivity to the interaction between climate processes in mid-latitudes and tropical latitudes (Giorgi & Lionello, 2008). Indeed, the reconstruction of the last 500 years’ climate indicates a negative winter rainfall trend since the 1960s in the Mediterranean region (Caloiero et al., 2018; Casas-Gómez et al., 2020), with higher reductions during the wet season (October-March; Esteban-Parra et al., 2022; Zittis et al., 2021). Since the end of the 20th century, climate change models predicted a further increase in temperatures and a decrease in precipitation, which has likely resulted in an increase in the frequency and intensity of extreme climate events such extreme droughts (IPCC, 2021; Lindner et al., 2010; Moemken & Pinto, 2022). This increase in drought stress has been considered the main abiotic cause of the observed dieback in Mediterranean forests (Camarero et al., 2015; Italiano et al., 2024). In particular, the Gibraltar strait region at SW Europe, which is notable for its rich biodiversity (Díaz-Villa et al., 2003), is experiencing a strong and consistent decline in forest cover (Jurado-Doña et al., 2022, Gutiérrez-Hernández & García, 2024).

Climate change modifies the distribution, physiological functioning and persistence of forest ecosystems. For instance, the increase of temperature has different implications such as the changes in the growing season length (Menzel & Fabian, 1999), compromising tree carbon stocks (Colangelo et al., 2021). Furthermore, southern Europe presents a negative trend of precipitation, being water availability, a critical factor limiting plant growth (Vicente-Serrano et al., 2014), commonly inducing tree physiological adjustments and feedbacks. For example, drought conditions will promote the decrease in stomatal conductivity, but negatively affect photosynthesis (Colangelo, et al., 2018; Muller et al., 2011; Palacio et al., 2014). Consequently, tree species under climate change conditions commonly experience growth reductions and/or defoliations, increasing the mortality risk (Sánchez-Salguero et al., 2012, 2013; 2020). It is therefore vital to gain a deeper understanding of the vulnerability of tree populations to extreme climate events, in order to predict and mitigate the adverse effects of climate change on Mediterranean forest ecosystems.

Among the multiple potential responses of tree species to drought, acclimation represents a crucial strategy for coping with new climatic conditions (Bose et al., 2024; Limousin et al., 2022). The acclimation of trees to drought events is influenced by a multitude of factors, including previous drought history (Vicente-Serrano et al., 2013), life history strategies, tree size, age, and microclimatic conditions related to topography, among other factors (Sánchez-Salguero et al., 2015; Anderegg et al., 2016; Bose et al., 2020; Gazol & Camarero, 2022). Furthermore, differences in growth resilience are associated with the loss of adaptive plasticity in drought events, which can be quantified by comparing the growth before and after the drought event (Lloret et al., 2011; Gazol et al., 2018). Some tree species show a strong growth resistance and low growth recovery, while others exhibit the opposite pattern (e.g., Anderegg et al., 2015; Bose et al., 2020; Sánchez-Salguero et al., 2018). Consequently, it is important to determine the capacity of tree species to cope with extreme events across environmental gradients to accurately assess their vulnerability in the context of a changing climate in the Mediterranean forests.

Oak species are among the most important in terms of economic, cultural, and ecological values in the Mediterranean region (Nixon, 2006; Plieninger et al., 2021). However, these species are facing an alarming rise in mortality across the Northern Hemisphere (Gentilesca et al., 2017). In Spain, this mortality phenomenon is commonly known as “*la seca*” and has been reported since 1990s (Brasier, 1992; Hernández-Lambraño et al., 2019). This dieback process affects different species of the *Quercus* genus such as *Q. ilex* L., *Q. suber* L. and *Q. canariensis* Willd., resulting in an increase in defoliation and subsequent tree mortality (Camarero et al., 2016; Hereş et al., 2018). Among these, *Q. canariensis* is an isohydric, ring-porous, and winter-deciduous species (Gea-Izquierdo et al., 2012; Sánchez-Salguero et al., 2020), with a high ecological relevance in Mediterranean forests, but highly susceptible to increasing aridity. However, there is a lack of knowledge regarding the ecological conditions that trigger the dieback process and the resilience of the species to extreme droughts.

In this study, we investigate the increasing dieback process of *Quercus canariensis* observed at Los Alcornocales Natural Park (southwestern Spain). We applied dendrochronological methods to characterize long-term growth responses to climate and extreme drought events across a sea-inland gradient differing in local microclimate. In this regard, we hypothesized that (1) forests that experienced higher aridity will show a decline in growth performance linked to a higher precipitation sensitivity, (2) due to the increasing trend in aridity, *Q. canariensis* will have a lower capacity to withstand extreme droughts in the recent decades.

## Materials and methods

### Field site and focal species

The study was performed at Los Alcornocales Natural Park, Spain (36° 25′ N, 5° 34′ W; Fig. 1). The area is mainly dominated by a mixed *Q. suber*-*Q. canariensis* forest, one of the most important Mediterranean oak woodlands, where several cases of forest dieback have been previously reported (Gómez-Aparicio et al., 2012; Gutiérrez-Hernández & García, 2024; Sánchez-Salguero et al., 2020). The predominant bedrock at this site is made by Oligo-Miocene siliceous sandstone soils, resulting in acid and sandy soils (Jurado-Doña et al., 2022; Paneque & Bellinfante, 1999). The mountains in the southern part of the Natural Park are oriented in a north-south direction, while in the north, they are oriented in a south-east orientation, with an elevation ranging from the sea level to 1092 m (Fig. 1). The climate in the study area is sub-humid Mediterranean, with dry summers and wet winters. The mean annual temperature is 15.7 °C, with January being the coldest month and July the warmest period. The mean annual rainfall for the period 1950-2020 is 589 mm year^-1^ (obtained from the closet meteorological station at the centre of the Natural Park, Cortes de la Frontera; 36° 32’ 40’’N, 5° 23’ 1’’W), although in the south, the dominant winds take humidity from the sea, creating a fog that is not recorded by the weather stations (Table 1; Jurado-Doña et al., 2022). This fog generates a microclimate that softens the summer drought in this part of the natural park.

**Figure 1.**
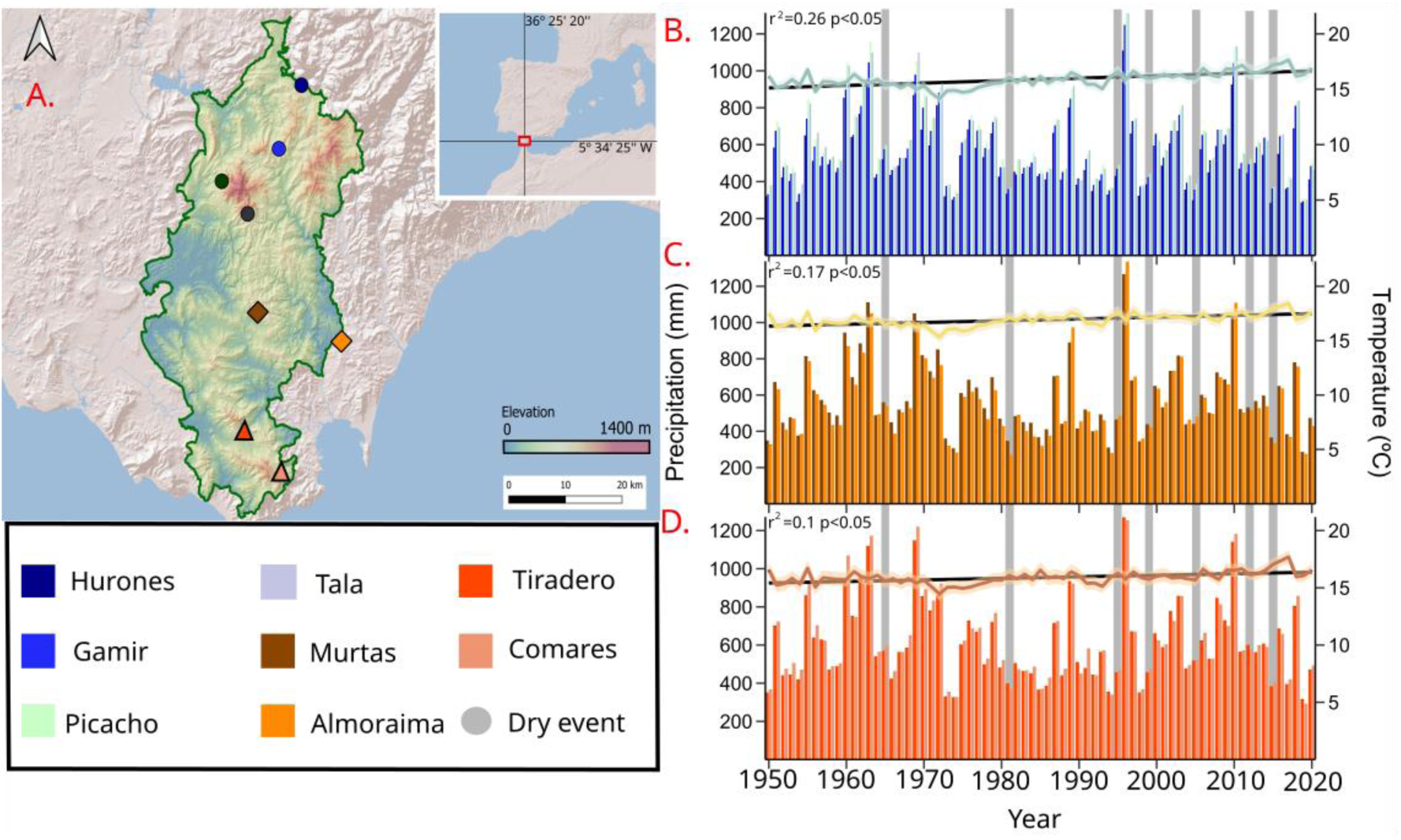
Location of the study sites within Los Alcornocales Natural Park with the 8 populations distributed in three different regions North (circles), Center (diamond) and South (triangles) (A). Annual precipitation (bars) at North (B), Center (C) and South (D) regions. The mean annual temperature since 1950 (smooth line) for each population is also displayed as well as the statistics their temperature trends (r^2^, correlation of determination and p, probability level). Each region is composed by different populations (identified by different colors, ranging from darkest (North) to lightest (South), following the latitudinal gradient. Grey areas in plots B, C and D indicate the selected drought events.

**Table 1.** Climatic, stand and tree characteristics of the three studied regions. Each region contains summarised information (mean and ±SD) of two or three sampled populations (see Field sampling section). Climate data (mean ±SD) values of maximum mean annual temperature (MMT), minimum mean annual temperature (mmt), and mean annual precipitation of the study sites for the period (1950-2020). Dendrochronology section describes stand and tree characteristics: mean values of tree density, number of sampled trees (and number of cores within brackets), time span of the tree-ring series, tree defoliation (%), diameter at breast height (DBH), tree height, tree minimum age at 1.3 m (age), mean basal area increment for the 1900-2021 period (BAI) and TRW first-order autocorrelation (AC). Different letters indicate significantly (*p*< 0.05) significant differences in mean values among regions based on Tukey tests.

| Site | North | Center | South |
| --- | --- | --- | --- |
| **Climate** |  |  |  |
| Elevation (m asl) | 460.8 ± 162.8 | 269.0 ± 147.0 | 337.0 ± 140.0 |
| MMT ($^{\circ}$C) | 20.4 ± 1.0a | 20.6 ± 0.7a | 19.1 ± 0.8b |
| mmt ($^{\circ}$C) | 11.35 ± 0.8a | 13.1 ± 0.9b | 12.6 ± 0.9c |
| Precipitation (mm) | 583 ± 207a | 574 ± 207a | 618 ± 218a |
| **Stand** |  |  |  |
| Tree density (ind. ha<sup>-1</sup>) | 231.1 ± 24.7 | 325.5 ± 213.1 | 220.0 ± 77.6 |
| N. Trees (cores) | 52 (97) | 29 (56) | 35 (63) |
| Time span (years) |  |  |  |
| **Tree** | 1744-2022 | 1778-2022 | 1856-2022 |
| Defoliation (%) | 50 ± 0.8a | 40 ± 0.6a | 50 ± 0.7a |
| DBH (cm) | 46.3 ± 15.8a | 50.7 ± 20.4a | 45.7 ± 18.3a |
| Height (m) | 15.8 ± 2.9a | 12.3 ± 2.6a | 18.3 ± 2.0a |
| Age (years) | 92 ± 45a | 79 ± 50ab | 71 ± 39b |
| BAI (cm<sup>2</sup> year<sup>-1</sup>) | 6.3 ± 3.1a | 7.9 ± 5.8b | 10.3 ± 5.1c |
| AC | 0.65 ± 0.15 | 0.53 ± 0.17 | 0.58 ± 0.23 |

The *Q. canariensis* is a deciduous tree native to southern Portugal, Spain, Tunisia, Algeria and Morocco. It is found in acid soils in the most humid parts of the valleys in the Mediterranean coastal area (Costa et al., 1998). This species tolerates a wide range of temperature conditions (0-24°C; Costa et al., 1998), although it is very sensitive to dry and warm spring conditions (Gea-Izquierdo et al., 2012). In the study area, *Q. canariensis* is associated with streams and valley bottoms, where the microclimate conditions are more humid (Urbieta et al., 2008). There are many plant species associated with this species, including the epiphytes *Polypodium cambricum* L. or *Davallia canariensis* L., as well as some vine-like species such as *Smilax aspera* L., *Hedera helix* L., *Lonicera periclymenum* L. or *Tamus communis* L. (Pérez-Ramos & Marañón, 2009).

### Field sampling

Eight different mixed-forest populations dominated by *Q. canariensis* were selected during 2022 along a latitudinal gradient within Los Alcornocales Natural Park, namely Hurones, Gamir, Picacho, Tala, Murtas, Almoraima, Tiradero and Comares, from north to south (see Fig. 1). Populations were selected within the protected area, trying to maintain similar levels of tree density, slope and exposition among populations (Table S1). In each population, 15 dominant adult trees were randomly selected, covering the natural size distribution of the forest. Populations were grouped in three regions within the latitudinal gradient defined by the distance to the sea (Fig. 1). The resulting categories were south (0-25 km away from the sea; Comares, Tiradero), center (26-50 km away from the see; Almoraima, Murtas) and north (51-75 km away from the see; Tala, Picacho, Gamir, Hurones).

Each tree was georeferenced using a GPS device (GPSMAP 66st, Garmin Ltd., USA), also recording the height of the tree using a clinometer, the diameter at breast height (DBH, measured at 1.3 m) and the tree health using the semi-quantitative scale proposed by the ICP Forest Network (Fischer & Lorenz, 2011): no defoliation (0%), light defoliation (1%-25%), moderate defoliation (26-60%), severe defoliation (61-99%) and dead (100%). To characterise the density of the populations, the position, DBH and species identity of each tree in the plot were recorded using a LiDAR sensor and the app ForestScanner (Tatsumi et al., 2022).

### Tree sampling and ring-width measurements

Two cores per tree were extracted at 1.3 m height using a 5-mm Pressler borer. Wood samples were air-dried, mounted and polished with successively finer grit sandpaper until the rings were visible (Fritts, 1976). Tree rings were first visually cross-dated, and tree-ring widths measured to the nearest 0.01 mm using a binocular microscope coupled to a computer with the LINTAB-TSAP^TM^ (Rinntech, Heidelberg, Germany). Then, tree-ring width series were evaluated and cross-dated per population using the COFECHA software (Holmes, 1983). The number of annual rings was counted to estimate age at breast height. Two different detrending approached were applied. First, to evaluate long-term growth trends, we accounted for the age and size long-term trends from the tree-ring width (TRW, mm year−^1^) series by converting them into basal area increments (BAI_t_, cm^2^ year^−1^), using the formula:

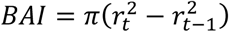

where *r* represents the tree radius increment and *t* indicates the year of the ring formation. BAI series were first averaged at tree level and then, at the population level to create the mean population BAI chronology. The BAI growth trends from 2000 **onward** were tested using linear regression models. Second, to evaluate climate-growth relationships, age- and size-related long-term trends were removed from the tree-ring width raw series applying a 30-year spline, thereby preserving high-frequency growth variability. The observed values were then divided by the fitted values to obtain dimensionless, standardized ring-width indices (Fritts, 1976). The resulting detrended series of ring-width indices (RWI) were combined to create a standard chronology for each population, using the biweight robust mean (Fig. S1). Detrending procedures were executed using the R package *dplR* (Bunn, 2008). Moreover, in order to establish a regional chronology, a principal component analysis (PCA) was conducted on the set of the population chronologies for each region, extracting the first component of the data analysis, explaining 58% of the variability of data from the North, 53% from the center and 45% from the southern region. This procedure aimed at extracting the common growth signal among populations of the same region.

### Climate-growth relationships and statistical analyses

The daily climate data series since 1950 were obtained from the interpolation of the three nearest meteorological stations using the R package *easyclimate* (Cruz-Alonso et al., 2023), this package get high-resolution daily climate data from European climatic database (E-OBS). Moreover, climate data comparisons of variables across populations were initially assessed for normality using Shapiro-Wilk tests. Subsequently, we used ANOVAs to compare climatic variables across populations, as all of the variables in question were parametric (Table S1; Bunn, 2008).

The climate-growth relationships were quantified for the period 1950-2020 using bootstrapped Pearson correlations with the *treeclim* R package (Zang & Biondi, 2015). These were estimated at population and regional level. The 1950-2020 period was selected due to the fact that 1950 represents the earliest year with reliable climatic data available from the study site, and it covered the majority of the population timespans (except for Comares; Table S1). Correlations were calculated among monthly mean temperature and total monthly precipitation data and population RWI chronologies. Seasonal correlations were also calculated (December previous year, January and February, winter; March, April and May, spring; June, July and August, summer; September, October and November, autumn) by averaging (temperature) and summing (precipitation). To estimate climate-growth relationships, correlations were calculated between regional chronology (obtained from PCAs) and regional climate data. For the regional climate data, we use the average monthly mean temperature and monthly precipitation of the populations in the region. Additionally, correlations were also calculated between the population chronologies and 1 to 20-months of Standardized Precipitation Evapotranspiration Index (SPEI) values (Fig. S2; Vicente-Serrano et al., 2010). In order to determine the correlation between regional growth and SPEI, we use bootstrapped correlations between regional chronologies and regional SPEI, using the regional climate data to obtain the regional SPEI value. The potential evapotranspiration (PET) was calculated using the Hargreaves approach (Droogers & Allen, 2002). The SPEI data were obtained using the *SPEI* R package (Vicente-Serrano et al., 2010).

### Resilience components

To evaluate the impacts of extreme droughts that occurred since 1950, we examined the 12 month-July SPEI values for the entire region (Fig. S3). We considered extreme drought years those with 12-month July SPEI lower than −1.0. Extreme drought years were then selected for the periods 1950-1999 and 2000-2020 (Sánchez-Salguero et al., 2018). We separated these two periods because i) previous studies have already highlighted a shift in the growth dynamics of deciduous oak species in the Mediterranean region from year 2000 (Colangelo et al., 2018; Sánchez-Salguero et al., 2020) and ii) there has been a general and significant increase in number of drought events in the southern Spain (González-Hidalgo et al., 2018a; Noguera et al., 2020a).

The components of resilience were defined based on BAI series following Lloret et al. (2011). The resistance, recovery and resilience indices were calculated using the following formulas:

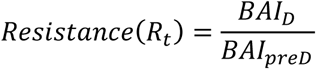

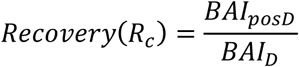

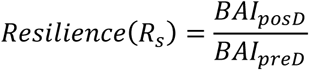

where BAI_D_ is the BAI during the corresponding drought year, BAI_preD_ is the average BAI for the 3 years preceding the drought event and BAI_postD_ is the average BAI for the 3 years following the drought event (Bose et al., 2020; Gazol et al., 2018; Pérez-Luque et al., 2021), except for the year 2021, where only 2 years were included due to the lack of post-drought ring-width data (Fig. S3). All these indices were calculated at the individual tree level for each of the drought periods studied.

To assess the capacity of the tree populations to withstand extreme droughts over time, we first considered the effect of the intensity of the drought events on the resilience components. To do this, linear models were constructed between the resilience components (Rs, Rc, and Rt) and the actual SPEI value for the given year (Table S2). Subsequently, the residual values of the model were extracted and modelled against time to illustrate the varying trend in resistance and recovery in both periods, as well as the resilience trend throughout the complete study period (given the lack of differences between them; Fig. S4). We also used ANOVAs to compare changes in Rt, Rs, and Rc between regions at every given event (Table S3). All the analyses were performed using R environment (version 4.3.1) and graphics with the R package *ggplot2* (Wickham, 2016).

## Results

### Climate trends and growth patterns

Climate data indicate that the warmest month is July, with an average temperature of 22.2 ± 1.1 °C, while the coldest month is January (9.0 ± 1.0 °C). The wettest and driest month are December (94.8 ± 87 mm) and July (0.4 ± 0.2 mm), respectively. However, the minimum and maximum annual temperatures exhibit a latitudinal gradient, with warmer temperatures at the northern sites and the southern sites have milder temperatures (Table 1). There has been a significant increase in temperature across all the populations since 1950, with a slope of 0.02 °C yr^-1^ (p < 0.05; Fig. 1B, C and D). Moreover, since 1950, we observed an increase in frequency of drought events, associated with low SPEI values (Fig. S3). No population showed a significant trend in annual precipitation since the 1950s. However, the central and northern regions have experienced the lowest annual precipitation records during the last decade (year 2019: 338.91 mm in the Center and 463.34 mm in the North).

We did not find significant differences in tree size, height and defoliation among populations of the latitudinal gradient (Table 1). Northern populations had older trees than southern ones, being Comares the youngest population and Gamir the oldest (Table S1). Consequently, the mean BAI for the period 1950-2021 was higher in the south, while the north presented the lowest mean BAI values (Table 1). Since 2000s, *Q. canariensis* growth exhibited similar increasing trends across the study populations and regions (Fig. 2), except for the center region.

**Figure 2.**
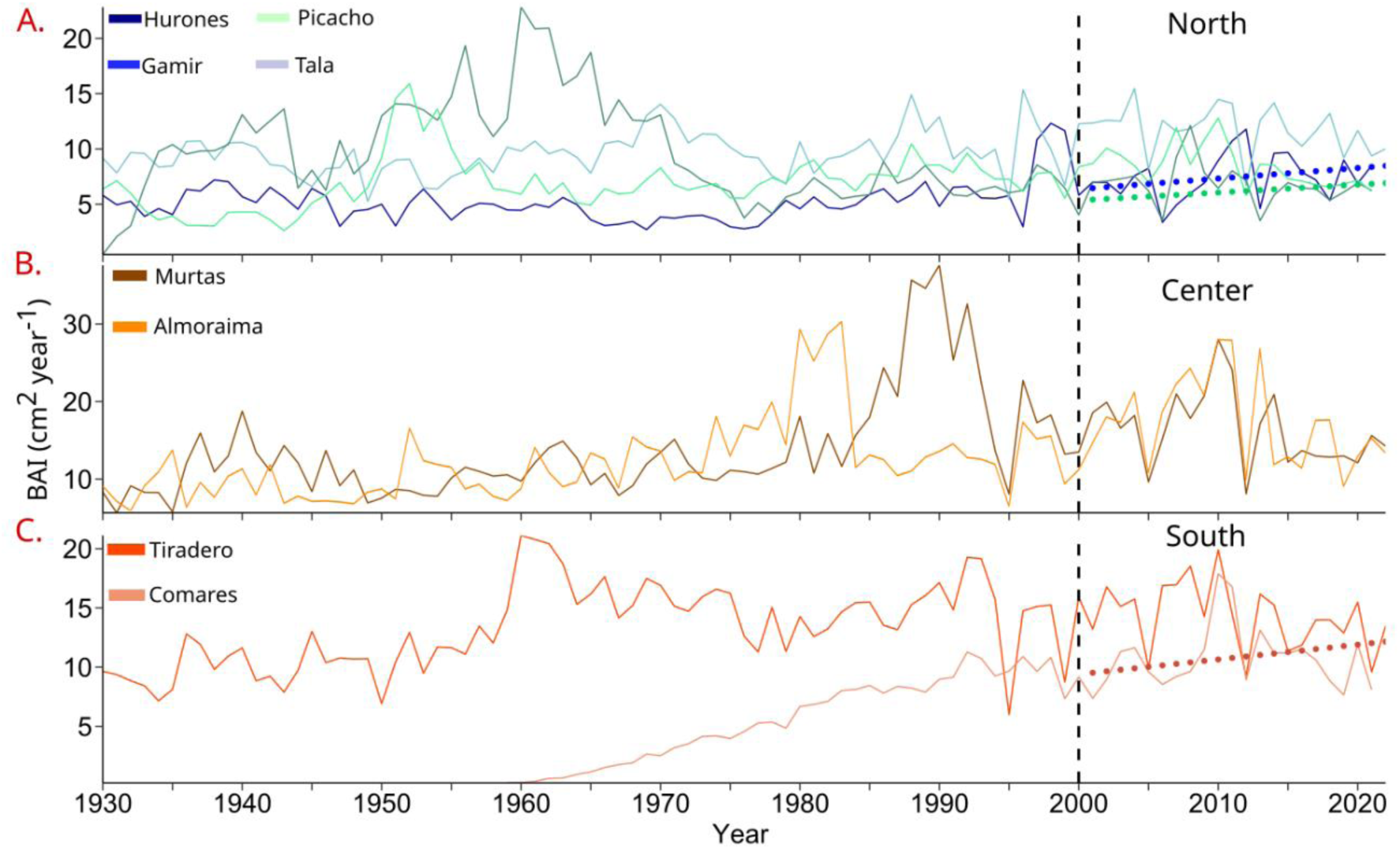
Basal area increment (BAI) series of the different study regions (A, North; B, Center; C, South). The vertical dashed lines indicate the year 2000. Dotted lines indicate significant growth trends (p< 0.05) since the year 2000 for each population.

### Climate-growth associations

Precipitation has the most significant influence on tree growth across all regions, though the temporal response window varies along the latitudinal gradient (Fig. 3). In the northern region, winter precipitation (December and January) and annual precipitation showed the strongest correlations. In the central and south region, this window is longer including previous august, late-winter and may precipitations (Fig. 3; Table 2).

**Figure 3.**
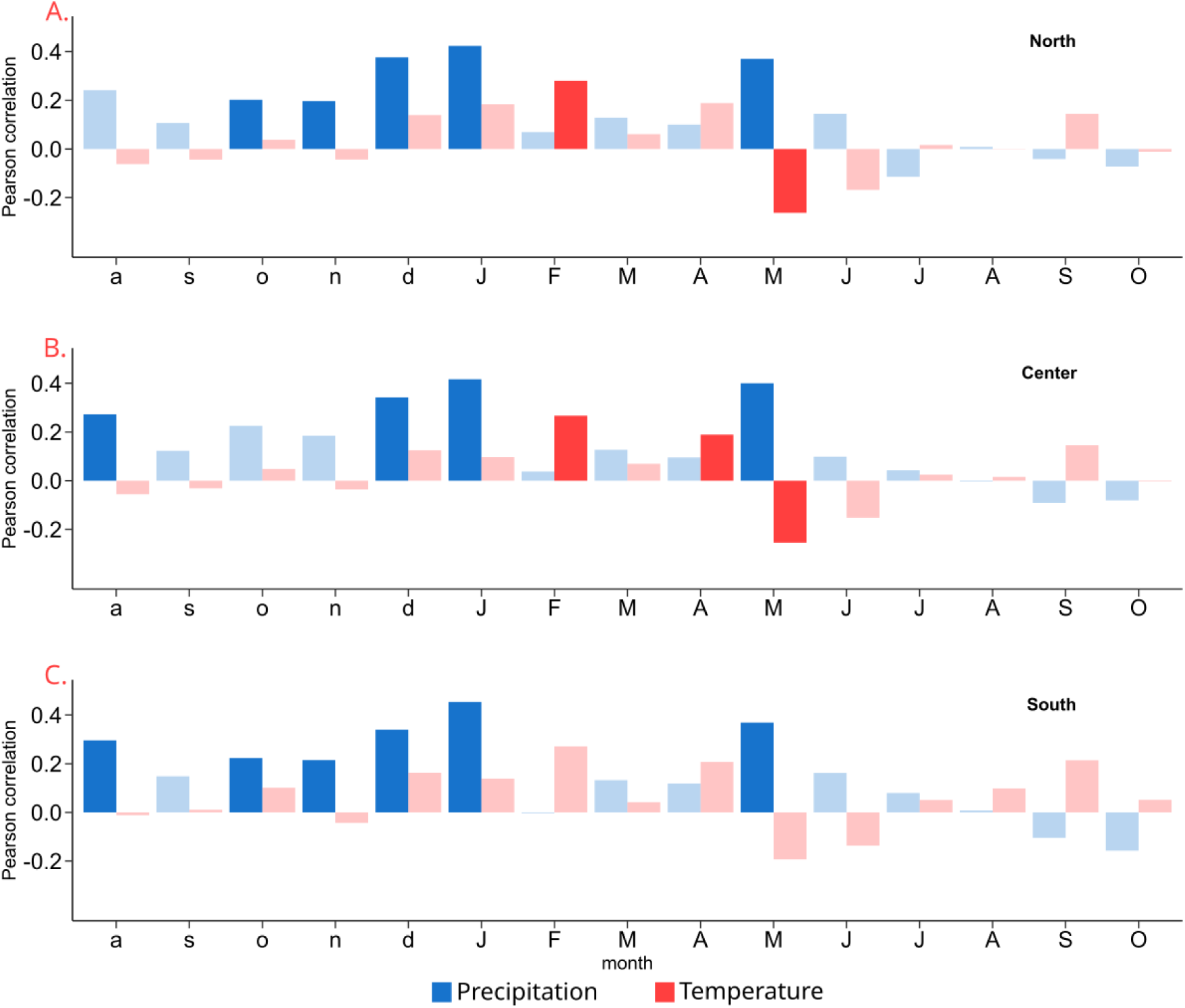
Bootstrapped Pearson correlations between growth and regional climate (temperature and precipitation) across the latitudinal gradient for the period 1950-2000. Lowercase letters indicate the month of the year preceding the ring formation, while uppercase letters indicate the months of the current year. Pale bars indicate non-significant correlations (*p*>0.05).

**Table 2.** Climate-growth relationships across regions along the latitudinal gradient. These are based on correlations between regional chronologies, seasonal, annual mean temperature and total precipitation of the regional climate for the period 1950-2000. Asterisk indicates the significance at p<0.05.

|  | Season | Site |  |  |
| --- | --- | --- | --- | --- |
|  |  | North | Center | South |
| Temperature | Year | 0.03 | 0.02 | 0.07 |
|  | Autumn<sub>p</sub> | -0.04 | 0.05 | 0.05 |
|  | Winter | 0.02 | 0.06 | 0.12 |
|  | Spring | 0.02 | 0.12 | 0.09 |
|  | Summer | -0.04 | 0.05 | 0.12 |
|  | Autumn | 0.03 | 0.00 | 0.03 |
| Precipitation | Year | 0.26 (*) | 0.39 (*) | 0.16 |
|  | Autumn<sub>p</sub> | 0.15 | 0.18 | 0.20 |
|  | Winter | 0.30 (*) | 0.48 (*) | 0.15 |
|  | Spring | 0.19 | 0.34 (*) | 0.30 (*) |
|  | Summer | 0.09 | 0.06 | 0.04 |
|  | Autumn | 0.05 | 0.11 | 0.02 |

We found few significant relationships between the growth of *Q. canariensis* and temperature across populations along the latitudinal gradient (Fig. 3). In the northern and central regions, February temperatures showed the strongest positive correlation with growth, while in contrast, May temperatures in the central region had a negative correlation. The south region did not show any significant correlation between growth and temperature. However, individual populations did show some significant relation between growth and temperature (Fig. S4). Tree growth of most of the northern populations showed a negative correlation with June temperature, while at the southern sites there is a significant correlation with warmer temperature during February (Tiradero) and September and tree growth (Comares; Fig. S4).

The SPEI exerted a significant effect on tree growth at short, mid and long-term time windows (1-20 months; Fig. 4, Fig. S2). Overall, tree growth of *Q. canariensis* showed significant associations to SPEI during the growing season (late-spring, summer and autumn). The populations most strongly associated with this drought index were those located at the north region, with a maximum correlation in late-winter drought (January-February) and spring-summer mild-term drought (r = 0.63; Fig. 4b). The southern region presented a lower correlation with SPEI, with the maximum correlation between May and September (8–15-month SPEI; Fig. 4c).

**Figure 4:**
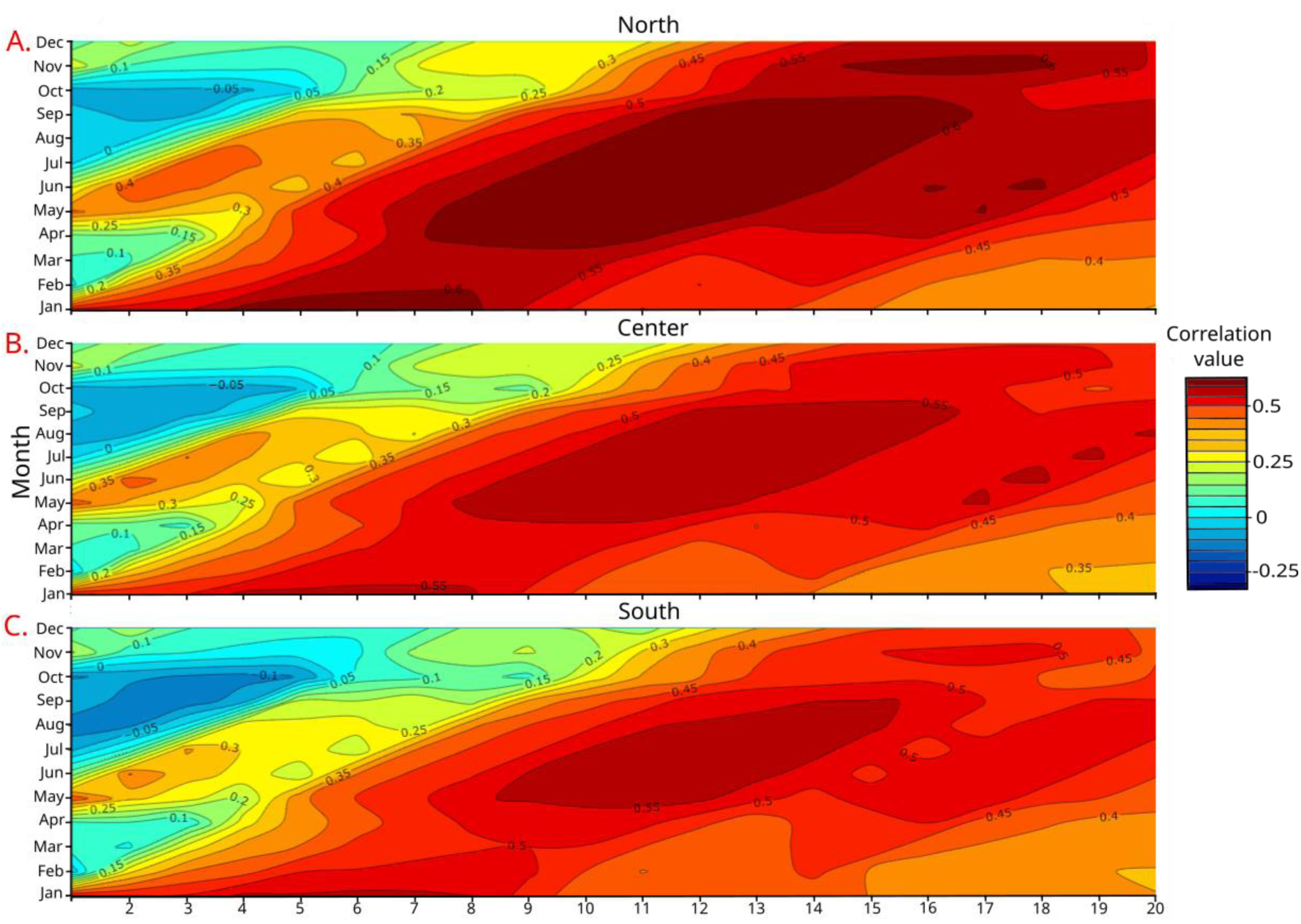
Bootstrapped Person correlations between regional chronologies and SPEI calculated at different time scales, accumulated from 1 to 20 months across the study regions.

### Resilience, resistance and recovery indices

The impact of extreme drought events on tree growth varied across the study regions. Resistance and recovery varied in *Q. canariensis* depending on the intensity of the drought event (Table S2), in contrast to resilience indices, which were independent from the intensity of the drought. Prior to 1999, there was no discernible trend in resistance, although a shift was observed from the 2005 drought. Between 2000 and 2020, resistance exhibited a general decline, with a consistent decline observed at the central region (Fig. 5 A, B) showing their minimum resistance value during the 2015 drought. In contrast, the recovery index displayed a positive trend throughout the 2000-2020 period, although the maximum recovery values were observed in the 1995 drought. These results also varied geographically across populations. The southern populations (Comares and Tiradero) exhibited a more pronounced decline in resistance than the populations situated at the centre of the gradient (Fig. S5). Moreover, since 2005 the only population that showed a negative trend in recovery was Picacho, which contrasted with the positive trend observed in other north population, Hurones. The growth resilience has been consistently declining at south region since the 1965 drought. Additionally, Tala population, in the north region, showed a stepper decline in this index since 1965 in comparison to Comares (Fig. S5 F). Finally, our findings showed similar resilience trends among regions during the different drought events.

**Figure 5.**
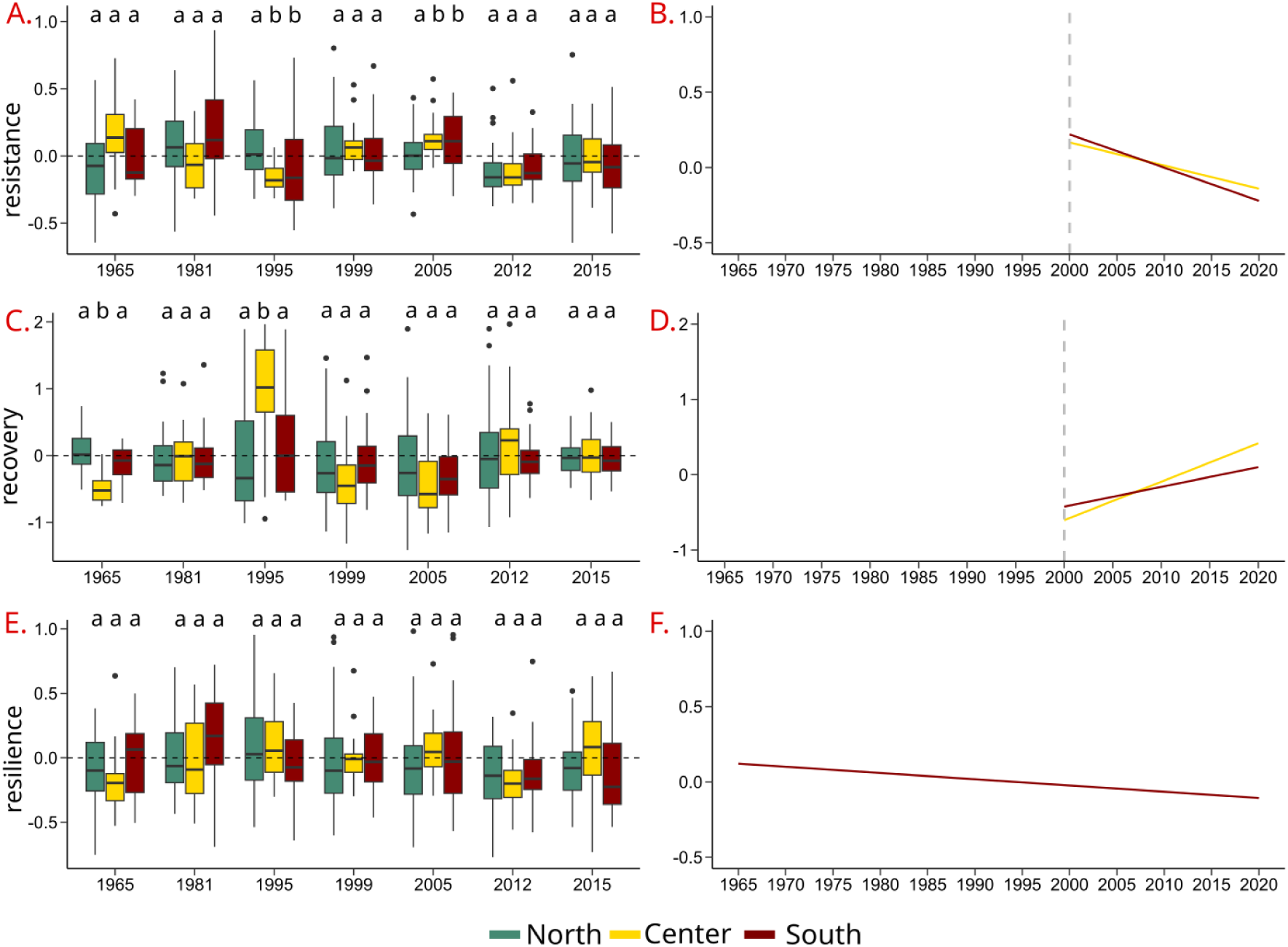
Resistance (A), recovery (C), and resilience (E) to extreme droughts during the 1950-2020 period calculated for individual tree BAI series and grouped by regions. These extreme droughts correspond to temperature and precipitation anomalies in 1965, 1981, 1995, 1999, 2005, 2012 and 2015 (Fig. S3). Significant trends (*p*< 0.05) of the respective indexes for the period 1965-2020 are shown in the panels B, D and F. When significant differences between the 1965-1999 and 2000-2015 period appeared, the results are presented separated in two blocks before and after year 2000 (vertical grey dashed line). Different letters indicate significantly (*p*< 0.05) different mean values across sites based on Tukey tests in each drought period.

## Discussion

In this study, we explored the growth performance and climate responses of an important western Mediterranean oak species over the last 75 years to assess the impact of climate change on its conservation. Our findings indicate that our study region has experienced an increase in temperature and in the frequency of drought events since the 1950s. Tree growth of *Q. canariensis* exhibits a differential sensitivity to climate across the study area, displaying grater sensibility to temperature and to mid-term drought at the north and center regions. However, tree growth of populations located at the south region is more sensitive to precipitation. This differential climatic sensitivity was also reflected in regional differences in the capacity to withstand extreme droughts. Populations in the southern regions exhibited changes in resistance and recovery, along with a significant decline in resilience over the past 75 years.

### Growth trends and spatial patterns of climate-growth responses

The study area exhibits a gradient of temperature that is influenced by its proximity to the sea (García-Mas et al., 2008). Our results demonstrate a consistent increase in temperature since 1950. There is no significant change in precipitation over the study period, although numerous studies predict a decline in precipitation for the coming decades (IPCC, 2021; Library et al., 2011). Our study reveals an increase in the frequency of drought events since the 2000s (González-Hidalgo et al., 2018b; Noguera et al., 2020b; Sánchez-Salguero et al., 2020). This increase in the frequency of severe drought events has significant implications for tree functioning (Camarero et al., 2016; Gazol & Camarero, 2022) and, at a broader scale, has already triggered numerous dieback episodes on different oak species (Colangelo et al., 2018; Fernández-Cancio et al., 2012; Sánchez-Salguero et al., 2020).

Except for Almoraina site, the wet sites with oceanic influence (south region) show higher growth rates in comparison to the other populations (Martínez-Vilalta et al., 2012; Morillas et al., 2024). This positive growth trend is usually related to young populations reflecting the relationship between vigour and longevity (Piovesan & Biondi, 2021). Instead, the population from driest region (center) shows a neutral growth trend over time (Fig. 2.B), despite being one of the youngest tree populations. In this case, we hypothesize that aridity exerts major effect on tree growth, which restrict *Quercus* growth even in the young individuals as also reported by Dhiedt et al. (2024). These results confirm *Q. canariensis* as a sensitive species to drought conditions, as occurs in others deciduous oaks in Mediterranean areas (Colangelo et al., 2021; Hernández-Lambraño et al., 2019).

Our results indicate that different climatic variables drive growth in *Q. canariensis* along the studied latitudinal gradient. Temperature has relatively a minor effect on the growth of *Q. canariensis*. However, the positive relation between growth and February temperatures could be linked to the reactivation of growth, finalising winter dormancy (Gea-Izquierdo et al., 2012). This fact occurs in trees in temperate climate regions when temperature reaches a certain value in spring (García González & Eckstein, 2003). In our study region, due to the mild winters, phenological observations indicate that cambial reactivation begins around March to April, depending on early spring and winter temperatures (Guada et al., 2021). Consequently, as a semi-deciduous oak, *Q. canariensis* exhibits a greater propensity for earlier leaf fall and earlier cambial reactivation in warmer early spring, which in turn results in the formation of larger vessels (Guada et al., 2020). Moreover, the center region and some northern populations show negative correlation between RWI and May temperature, which suggest that the stored sugars are consumed by the mid-growing season due to the heat stress (Gea-Izquierdo et al., 2012). This negative relationship is not evident at the southern region probably likely due to the milder temperatures experienced by populations closer to the sea.

Precipitation exerts a stronger control over growth than temperature at our study sites. This result is consistent with previous studies analyzing climate-growth relationships in this species (Sánchez-Salguero et al., 2020), although the temporal window differs. Our findings indicate that higher precipitation enhances growth, though there is some variability in seasonality across the studied latitudinal gradient. Furthermore, along this gradient, *Q. canariensis* growth is positively associated with wetter previous winters and the month of May in the current year. The impact of winter precipitation suggests that groundwater replenishment is crucial for this species, particularly given the sandy and shallow soil characteristics, which limit water retention capacity (Helama et al., 2009).

Several studies have identified that drought in springs and summers as the primary drivers of oak growth decline (Mausolf et al., 2018; Popa et al., 2013). Our results confirm that *Q. canariensis* is highly sensitive to drought between spring and summer, as occurs in other deciduous oaks (Petritan et al., 2021). In fact, growth of *Q. canariensis* presents the highest values of response to SPEI during the late-spring to summer period. This could be attributed to the fact that spring-deciduous species are more affected by dry springs, disturbing their cambial activity and altering the formation of new functional xylem vessels (Puchałka et al., 2024). If the drought persists throughout the summer season, trees initiate an early leaf senescence that limits carbon assimilation during the autumn (Pataki & Oren, 2003). Consequently, the limited water storage in spring and summer causes a severe reduction in latewood production in oak species (Alla & Camarero, 2012). We observe a gradient of responses across regions in the correlation between SPEI and growth, with the north and center regions showing the strongest correlations and south the weakest. These findings confirm the sensitivity of *Q. canariensis* to changes in climate and suggest that drier and warmer conditions may result in reduced growth, as has been observed in other deciduous oaks in Mediterranean areas (Gea-Izquierdo et al., 2012; Gentilesca et al., 2017). The central region, notable for its aridity, is identified in our analyses as the area where growth is the most susceptible to drought conditions since it did not show a positive growth trend, which can be attributed to the long-term unfavourable climatic conditions.

### Changes in resistance, recovery and resilience

We analysed the resistance, recovery and resilience to drought across an environmental gradient, considering seven extreme drought events registered between 1950 and 2020. Our demonstrate that resistance and recovery depend on drought intensity, which suggested that the physiological responses to drought increased with drought severity (Gao et al., 2018), showing a greater capacity to resist as more intense the drought is. We hypothesize that *Q. canariensis* has the potential to have a full growth recover due to its ring-porous water system, which allows for the rapid recuperation from drought conditions and the development of new small-size xylem tissues (Zweifel & Sterck, 2018). Therefore, under a future context of reduced precipitation and increased frequency and intensity of hotter droughts, the rapid growth recovery (three years) observed in the drier sites could be a pivotal factor governing the functioning of *Q. canariensis* forests.

There is a growing body of literature that describes and quantifies the interacting effects of drought and tree growth (Colangelo et al., 2021; Gazol et al., 2018; Martínez-Vilalta et al., 2012; Martínez-Vilalta & Piñol, 2002; Sánchez-Salguero et al., 2018, 2020). A shift in the resistance and recovery trends has been observed in the year 2000, suggesting a change in the growth strategy associated with the increased frequency of extreme drought events (Colangelo et al., 2021). The current strategy is based on a plastic response in xylem vessels to precipitation variability (Boseet al., 2024) and probably *Q. canariensis* may rebalance root and shoot biomass allowing for a full recovery after severe drought events (Móricz et al., 2021). However, this plastic response induces a legacy effect in tree growth, as shown in other oaks (Boseet al., 2024; Gazol et al., 2020). For example, this use of carbon resources for the recovery following drought could be a contributing factor to the observed decline in resistance, potentially due to the inability to replenish these resources at an increasing frequency of drought episodes, in line with results from other tree species (Kannenberg et al., 2019; Serra-Maluquer et al., 2018), being this trend more notable in cooler areas (south region; DeSoto et al., 2020).

*Q. canariensis*, like other trees in the Mediterranean region, shows a consistent decline in resilience to extreme drought events since 1950 at the coolest region (Gentilesca et al., 2017; Serra-Maluquer et al., 2018), commonly related with the hydraulic failure mechanism. This results in xylem embolism, which impairs the transport of water from the roots to the leaves (Colangelo et al., 2017). At the tree level, it leads to a canopy dieback and branch mortality (Adams et al., 2013), inducing a decline in resilience. In addition, it was observed that resilience did not demonstrate any trend in xeric regions, suggesting that populations from the north and central regions have adapted to those conditions making them more tolerant than their wetter counterparts and facilitate a more rapid recovery of growth following an extreme drought (Anderegg & Hillerislambers, 2016; Stuart-Haëntjens et al., 2018). In contrast, numerous studies have demonstrated that regions with cooler and wetter climates are more susceptible to extreme droughts (DeSoto et al., 2020; Gazol et al., 2018; Pardos et al., 2021). Thus, the observed decreased resilience trend in the south may be attributed to higher temperatures in the wet season, which intensify the drought effect, causing an alteration on the tree physiology (Gori et al., 2023). Moreover, ecophysiological processes exert an influence on tree growth and their resilience capactity, suggesting that resilience to drought may vary with individual size and age. Consequently, it may be necessary to incorporate additional factors in future resilience analyses, as other authors have reported (Castagneri et al., 2022; Merlin et al., 2015).

These results highlight the need of considering a range of spatial and temporal scales when studying forest responses to environmental alterations. The long-term variation in forest functionality is determined by multiple factors, thereby underscoring the importance of this approach to show the warning signals of oak decline in response to climate change (Lloret et al., 2011; Sánchez-Salguero et al., 2020). Our results also demonstrate that the populations in the southern and in the central regions are the most vulnerable to future climate, with the southern region exhibiting a negative resilience trend. *Q. canariensis* at the central region currently have a recovery capacity that compensates for the decline in growth following the drought event. Moreover, the current growth trend and its high-drought sensitivity indicate that this region is one of the most susceptible, showing early-signals of growth decline with the shift in resistance and recovery trend since 2000s. Our results show that the cumulative impact of drought impairs the resilience of *Q. canariensis* to cope extreme drought events, potentially threatening the long-term persistence of this species.

## Conclusions

The latitudinal pattern of climate conditions at the study area has resulted in differential growth trends of *Q. canariensis* among regions and populations. The main common environmental factor influencing the growth of *Q. canariensis* was winter precipitation. Drought exerted a significant role in *Q. canariensis* growth, particularly in the north and central region. In these regions, *Q. canariensis* may have some adaptations to resist drought events, although under a future drier scenario, these regions have been identified as being at risk of exceeding their capacity for drought resistance. Moreover, the central region shows an early signal of declining with a shift in resistance and recovery trend since 2000. However, no significant trend of growth decline has been found in the drier sites due to their drought sensitivity, negating our first hypothesis. In addition, the resilience of this species to extreme drought events has been in consistent decline in the southern region since the 1950s, confirming our second hypothesis. As a result, the most humid region is the most endangered with a lower capacity to withstand extreme drought events. Our approach enabled us to identify the vulnerability of this species to climate change by calculating its resilience to severe drought events, indicating a decline in this Mediterranean deciduous oak under an aridity increase scenario.

