## supplementary material for "Negative resilience trends to more frequent extreme droughts in Mediterranean oak species"

### Appendix

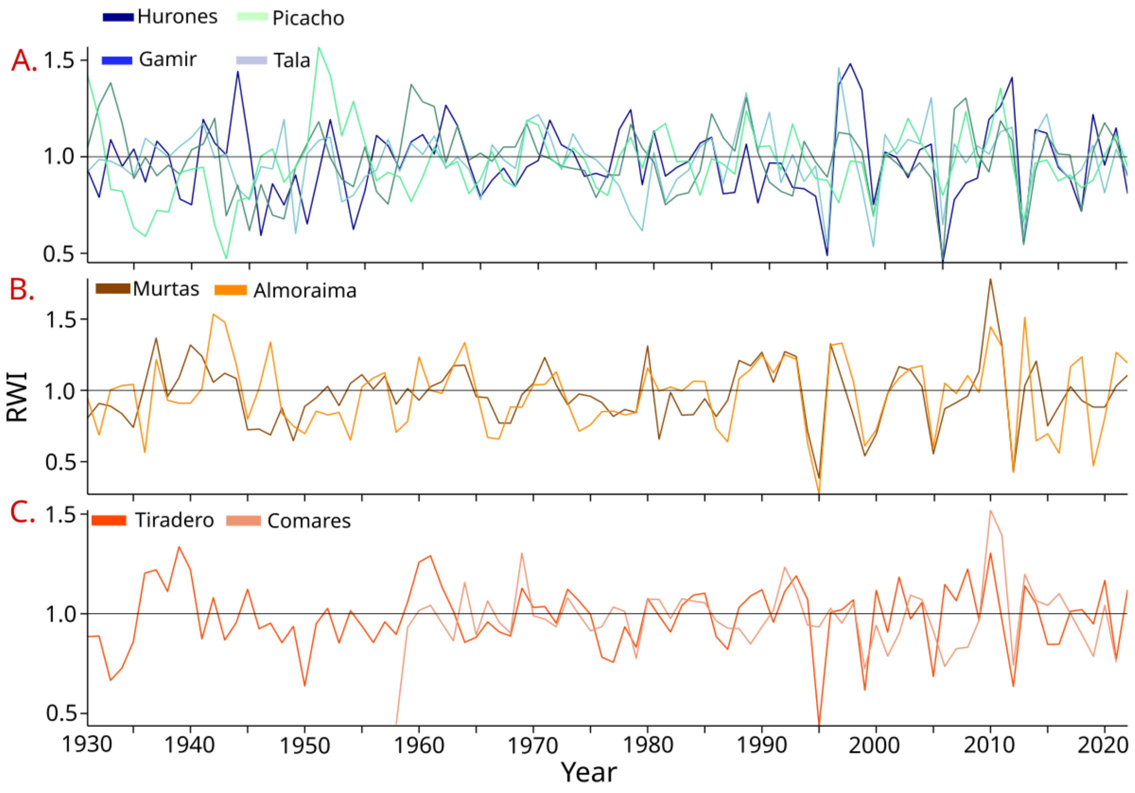


**Figure S1:** Standardized tree ring-width indices (RWI) series of the different study populations aggregated by regions A), North, B) Center, C) South.

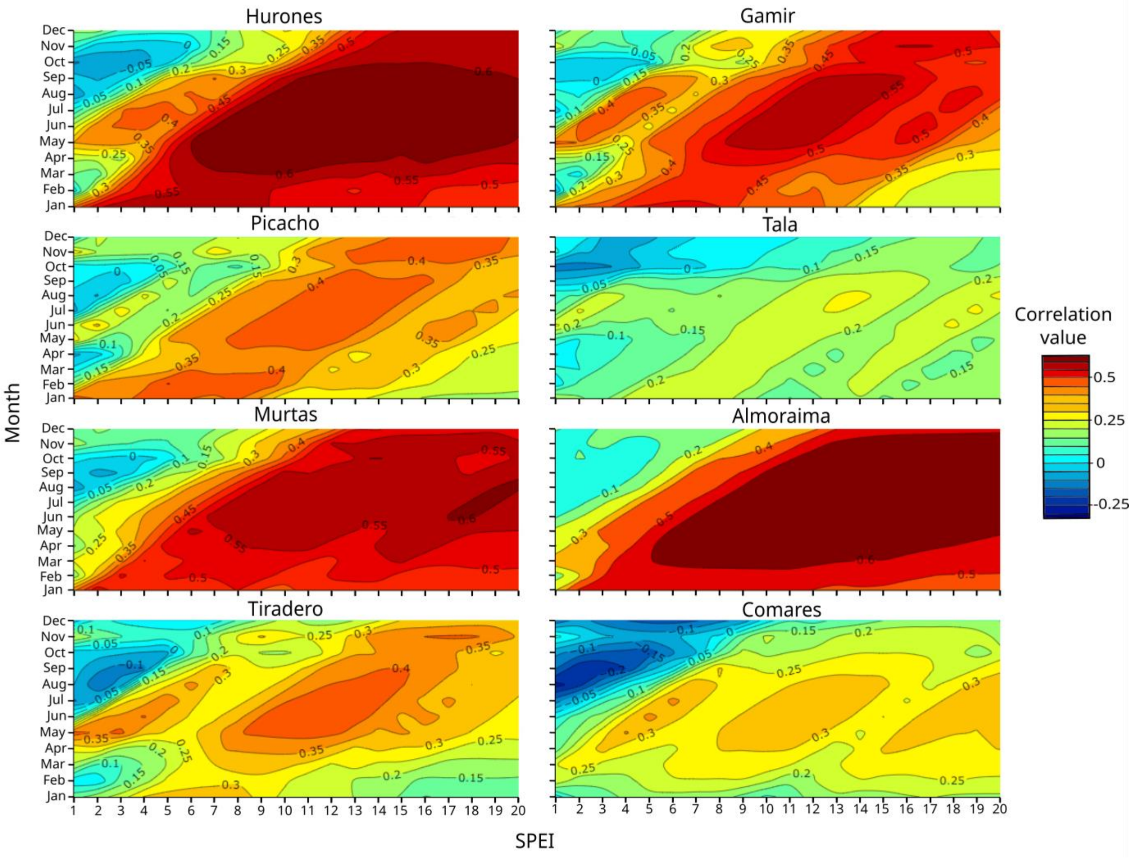


**Figure S2:** Relationships between the Standardized precipitation-evaporation index (SPEI) and growth at the different study populations based on Pearson correlations calculated by ring-width chronologies and monthly values (y axes) of the SPEI drought index calculated at several time scales, from 1 to 20 months.

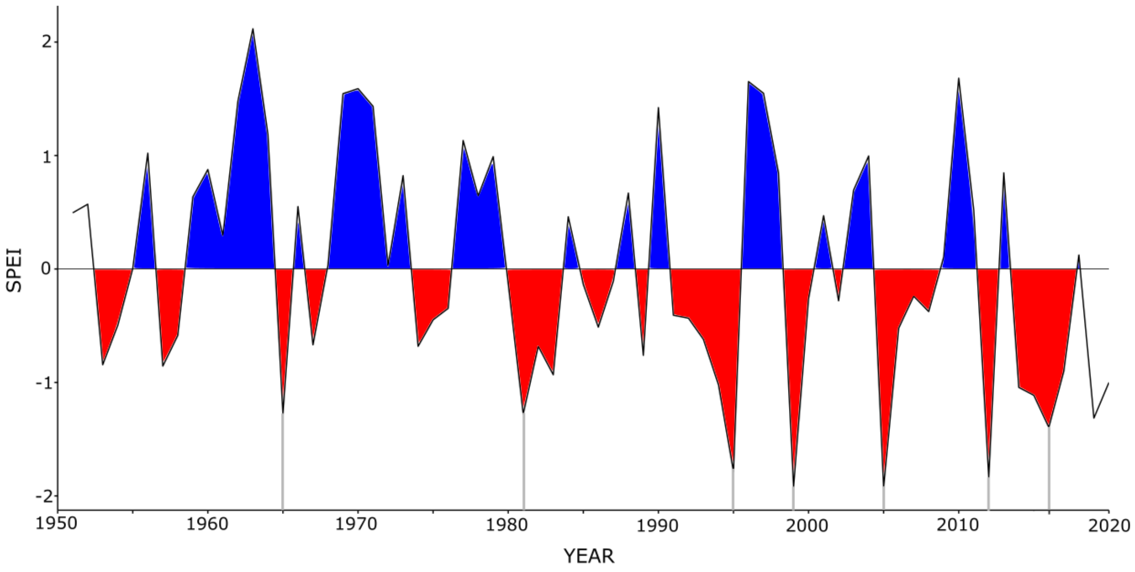

**Figure S3.** Standardized precipitation-evaporation index (SPEI) indicating drought severity values calculated at 12-month long scale from July for the 1950-2020 period. Blue areas show the wet periods, while red areas indicate the drought events. Grey lines indicate extreme drought years.

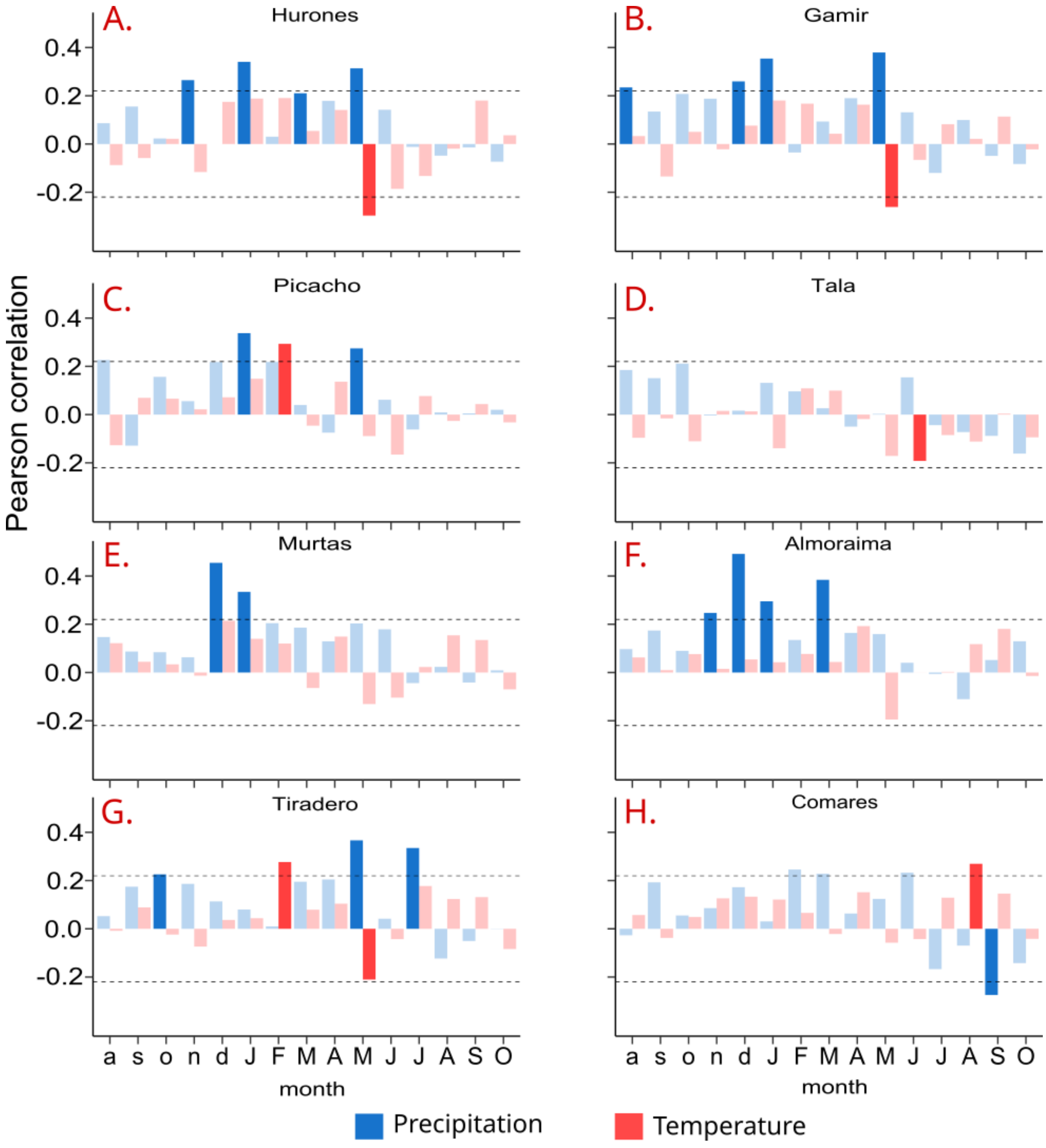


**Figure S4:** Climate-growth relationships at the different study populations. Bars correspond to Spearman correlations obtained by relating population mean RWI and monthly mean temperature (red) and total precipitation (blue) for the period 1950-2020. Dashed horizontal lines show the significance at p< 0.05 levels. Lower case letters indicate the months of the year preceding the ring formation, while capital letters indicate the months of the current year. The smooth bars indicate not significant correlation.

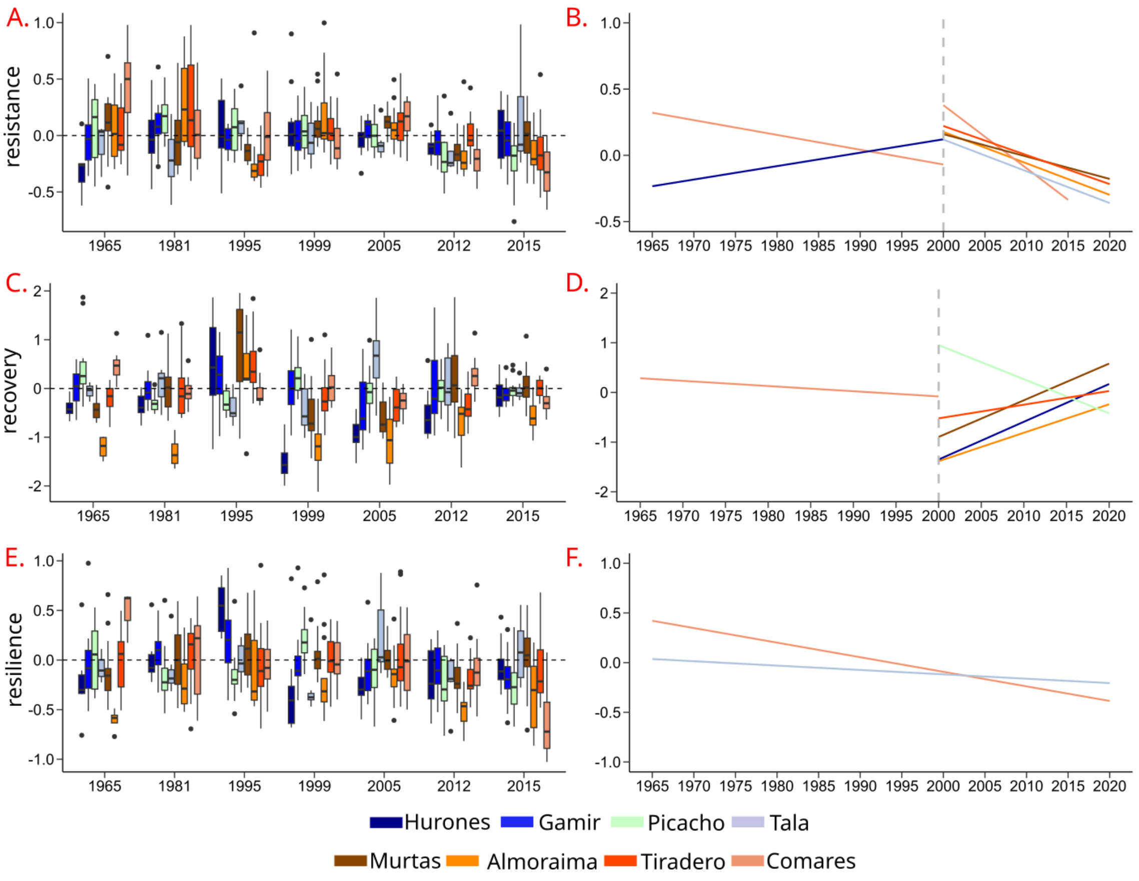


**Figure S5.** Resilience components, resistance (A,B), recovery (C,D), resilience (E,F), calculated for individual tree BAI series per site and extreme droughts during the 1950-2015 period. These extreme droughts correspond to temperature and precipitation anomalies in 1965, 1981, 1995, 1999, 2005, 2012 and 2015. Significant trends ($p < 0.05$) of the indexes for the period 1965-2020 are show in the panels B,D and F. When significant differences between the 1965-1999 and 2000-2015 period appeared, the results are presented separated in two blocks before and after year 2000. The dashed line in the panels A,C,E, represent the threshold of resistance, recovery and resilience index.

**Table S1.** Location and climate data (mean ±SD) values of maximum mean temperature (MMT), minimum mean temperature (mmt), and mean annual precipitation of the study sites. Dendrochronology section describes tree characteristics: mean values of density, number of trees and cores, recorded period, diameter at breast height (DBH), tree height, tree age at 1.3 m, mean basal area increment for the 1900-2021 period (BAI) and first-order autocorrelation (AC). Different letters indicate significantly ($p < 0.05$) different mean values across sites based on Tukey tests.

| site | Hurones | Gamir | Picacho | Tala | Murtas | Almoraima | Tiradero | Comares |
|---|---|---|---|---|---|---|---|---|
| **Climate** | | | | | | | | |
| Latitude | 36.66 | 36.56 | 36.51 | 36.47 | 36.33 | 36.29 | 36.15 | 36.10 |
| Longitude | -5.49 | -5.53 | -5.64 | -5.59 | -5.56 | -5.39 | -5.58 | -5.51 |
| Altitude (m a.s.l.) | 217 | 511 | 445 | 670 | 416 | 122 | 197 | 477 |
| Orientation | NE | SW | NW | NE | NE | NW | NE | NW |
| MMT (ºC) | 21.47 ± 0.95 a | 20.43 ± 0.73b | 20.19 ± 0.79bc | 19.90 ± 0.77cd | 20.26 ± 0.65bc | 21.12 ± 0.63a | 19.59 ± 0.66d | 18.72 ± 0.73e |
| mmt (ºC) | 12.11 ± 0.80d | 11.07 ± 0.69e | 10.98 ± 0.62e | 11.23 ± 0.62e | 12.54 ± 0.60c | 13.82 ± 0.68a | 13.43 ± 4.22b | 11.44 ± 0.60d |
| Precipitation (mm) | 524 ± 174a | 578 ± 194a | 616 ± 212a | 612 ± 211a | 585 ± 204a | 563 ± 210a | 610 ± 215a | 626 ± 221a |
| **Dendroconology** | | | | | | | | |
| Tree density (tres $ha^{-1}$) | 180.2 ± 40.1 | 231.1 ± 100.8 | 86.4 ± 60.6 | 301.0 ± 36.4 | 169.7 ± 57.8 | 248.2 ± 84.3 | 129.1 ± 21.7 | 562.9 ± 74.4 |
| No. Trees (cores) | 12(22) | 15(29) | 10(16) | 15(30) | 15(29) | 14(27) | 19(34) | 16(29) |
| Time spam | 1842-2022 | 1744-2022 | 1844-2021 | 1835-2021 | 1778-2022 | 1879-2022 | 1856-2022 | 1958-2021 |
| Defoliation (%) | 50 ± 60abcd | 70 ± 80 abcd | 1 ± 1ab | 10 ± 30bcd | 70 ± 60abc | 10 ± 30cd | 90 ± 80a | 0 ± 0d |
| DBH (cm) | 41.0 ± 14.5ab | 57.3 ± 12.8a | 38.9 ± 14.5ab | 42.5 ± 13.3ab | 53.7 ± 17.0a | 47.4 ± 22.3ab | 55.8 ± 18.1a | 34.0 ± 3.0b |
| Height (m) | 8.7 ± 1.1d | 14.0 ± 1.8a | 8.1 ± 1.2d | 12.7 ± 1.5ab | 12.2 ± 3.1abc | 12.4 ± 1.8abc | 10.4 ± 2.1cd | 11.4 ± 1.1bc |
| Age (yr) | 75 ± 38bc | 124 ± 41a | 76 ± 44bc | 80 ± 38bc | 103 ± 49ab | 53 ± 37c | 87 ± 47b | 52 ± 9c |
| BAI (cm2 year-1) | 5.8 ± 2.7c | 7.5 ± 3.2b | 4.5 ± 3.5d | 6.4 ± 2.3bc | 6.4 ± 5.8c | 10.4 ± 4.9a | 11.4 ± 5.0a | 7.4 ± 4.2bc |
| AC | 0.59 ± 0.17 | 0.66 ± 0.10 | 0.70 ± 0.13 | 0.67 ± 0.18 | 0.62 ± 0.14 | 0.43 ± 0.15 | 0.52 ± 0.27 | 0.66 ± 0.15 |

**Table S2.** F-value and P-Value of the Linal model between population or regional SPEI and resistance, recovery and resilience along 1965, 1981, 1995, 1999, 2005, 2012 and 2015 extreme drought events. Bolt numbers indicate p-value < 0.05.

| | Resistance | | Recovery | | Resilience | |
|---|---|---|---|---|---|---|
| | **F-Value** | **P-Value** | **F-Value** | **P-Value** | **F-Value** | **P-Value** |
| **Population** | | | | | | |
| Hurones | 36.77 | **<0.001** | 4.644 | 0.0346 | 0.42 | 0.516 |
| Gamir | 116.6 | **<0.001** | 49.44 | **<0.001** | 1.13 | 0.288 |
| Tala | 32.57 | **<0.001** | 10.13 | **0.001** | 0.96 | 0.328 |
| Picacho | 9.27 | **0.003** | 18.62 | **<0.001** | 0.94 | 0.336 |
| Murtas | 58.27 | **<0.001** | 24.14 | **<0.001** | 1.52 | 0.219 |
| Almoraima | 15.05 | **<0.001** | 2.10 | 0.151 | 0.42 | 0.518 |
| Tiradero | 49.78 | **<0.001** | 31.26 | **<0.001** | 0.30 | 0.579 |
| Comares | 34.62 | **<0.001** | 0.01 | 0.916 | 21.3 | **<0.001** |
| **Region** | | | | | | |
| North | 156.8 | **<0.001** | 80.69 | **<0.001** | 0.12 | 0.722 |
| Center | 78.75 | **<0.001** | 43.15 | **<0.001** | 2.05 | 0.153 |
| South | 61.2 | **<0.001** | 19.91 | **<0.001** | 4.263 | 0.060 |

**Table S3.** Differences in resistance, recovery and resilience between populations in 1965,1981,1995,1999,2005,2012 and 2015 drought, in base on Tukey test analysis. Different letters indicate significantly ($p < 0.05$) different mean values across sites based on Tukey tests.

| | Hurones | Gamir | Picacho | Tala | Murtas | Almoraima | Tiradero | Comares |
|---|---|---|---|---|---|---|---|---|
| **1965** | | | | | | | | |
| Resistance | a | ab | ab | ab | ab | ab | ab | b |
| Recovery | a | ab | ab | b | a | c | a | b |
| Resilience | ab | ab | ab | b | ab | a | ab | b |
| **1981** | | | | | | | | |
| Resistance | a | a | a | a | a | a | a | a |
| Recovery | a | a | a | a | a | b | a | a |
| Resilience | a | a | a | a | a | a | a | a |
| **1995** | | | | | | | | |
| Resistance | a | ab | a | a | ab | ab | b | a |
| Recovery | ab | ab | bc | b | a | ab | ac | bc |
| Resilience | a | ab | ab | b | ab | b | b | b |
| **1999** | | | | | | | | |
| Resistance | a | a | a | a | a | a | a | a |
| Recovery | a | bc | b | bc | c | a | bc | bc |
| Resilience | a | ab | ac | b | bc | ac | ab | ab |
| **2005** | | | | | | | | |
| Resistance | a | ab | a | ab | ab | ab | ab | b |
| Recovery | ab | acd | ce | ce | bd | b | cd | cd |
| Resilience | a | ab | b | ab | ab | a | b | b |
| **2012** | | | | | | | | |
| Resistance | ab | ab | ab | b | ab | ab | a | b |
| Recovery | ab | acd | bd | bd | cd | b | bc | d |
| Resilience | ab | b | ab | ab | b | a | b | b |
| **2015** | | | | | | | | |
| Resistance | ab | bc | b | ac | ab | bc | bc | c |
| Recovery | ab | ab | ab | ab | a | c | ab | bc |
| Resilience | ab | ab | bc | ab | b | ac | ab | c |
